# Multivariate associations between dopamine receptor availability and risky investment decision making across adulthood

**DOI:** 10.1101/2021.06.15.448537

**Authors:** Mikella A. Green, Kendra L. Seaman, Jennifer L. Crawford, Camelia M. Kuhnen, Gregory R. Samanez-Larkin

## Abstract

Pharmacological manipulations have revealed that enhancing dopamine increases financial risk taking across adulthood. However, it is unclear whether baseline individual differences in dopamine function, assessed using PET imaging, are related to performance on risky financial decision making tasks. Here, thirty-five healthy adults completed an incentive-compatible learning-based risky investment decision task and a PET scan at rest using [11C]FLB457 to assess dopamine D2-like receptor availability. In the task, participants made choices between a safe asset (bond) and a risky asset (stock) with either an expected value less than the bond (“bad stock”) or expected value greater than the bond (“good stock”). Five measures of behavioral performance (choice inflexibility, risk seeking, suboptimal investment) and beliefs (absolute error, optimism) were extracted from the task data and average non-displaceable dopamine D2-like binding potential was extracted from four brain regions of interest (midbrain, amygdala, anterior cingulate, insula) from the PET imaging data. Given the presence of multiple independent and dependent variables, we used canonical correlation analysis (CCA) to evaluate multivariate associations between learning-based decision making and dopamine function controlling for age. Decomposition of the first dimension (r = .76) revealed that the strongest associations were between measures of choice inflexibility, incorrect choice, optimism, amygdala binding potential, and age. Follow-up univariate analyses revealed that amygdala binding potential and age were both independently associated with choice inflexibility. The findings reveal latent associations between baseline neural and behavioral measures suggesting that individual differences in dopamine function may be associated with learning-based financial risk taking in healthy adults.

## Introduction

Why do people take risks? There are many potential contributions to risk preferences from biases in perceptions of risk, beliefs about benefits, differential weighting of costs and benefits, misunderstanding of risk or low numeracy in general, among many other factors (Hogarth, 1975; Kahneman & Tversky, 2012; Sjöberg, 2000). Some have suggested that the source of risk preferences may be at least partially based on prior learning (Hertwig, Barron, Weber, & Erev, 2004; Hertwig & Erev, 2009; Kuhnen, 2015). Many risks taken in everyday life rely on feedback from prior experience. Individuals’ predictions of the costs and benefits of making a specific choice may be initially biased but should be updated after experience realizing outcomes of choices. However, individuals vary in their rates of learning or willingness or ability to update expectations at all (Schönberg, Daw, Joel, & O’Doherty, 2007). Given how strongly dopamine has been implicated in updating expectations and learning in general (Schultz, 2002; Schultz, Dayan, & Montague, 1997), potential dopaminergic contributions to risk taking may be related to how organisms learn about risk. Individual differences in dopamine function may be related to learning-based financial risk taking in general.

Pharmacological modulation of the dopamine system has been shown to increase financial risk taking in everyday life for some individuals. The most well known examples are that some middle-aged and older individuals with Parkinson’s Disease (PD) who are treated with dopamine agonists develop novel gambling addictions and engage in other excessively risky behaviors (Weintraub et al., 2006). Relatedly, drugs that act on dopamine transporters are among the most commonly prescribed drugs to reduce impulsivity in children, adolescents, and adults (Piper et al., 2018). In children with ADHD, methylphenidate has been shown to decrease risk taking (DeVito et al., 2008). The same drugs have been shown to reduce excessive risk taking in older adults with frontal temporal dementia (FTD) (Rahman et al., 2006). The differences in the direction of these dopaminergic effects are unlikely to be due to differences in the mechanism of action of the drugs. Both agonists and transporter blockers increase dopamine levels in the synapse. These same groups of individuals with PD, ADHD, and FTD have also been shown to have baseline differences in dopamine function. Thus, individual differences in baseline dopamine levels may be directly associated with risk taking. In fact, studies with early-stage Parkinson’s patients suggest that baseline dopamine measures may predict adverse dopamine treatment responses that induce excessive risk taking (Steeves et al., 2009; Vriend et al., 2014). However, few studies have examined how individual differences in dopamine function are related to risk taking in general.

Both learning abilities and financial risk-taking vary across individuals of different ages. Studies of aging and reinforcement learning show that older adults learn more slowly (Eppinger, Hämmerer, & Li, 2011; Mell et al., 2005), but studies of aging and risk taking are not as clear (Rolison, Hanoch, Wood, & Liu, 2014). Meta-analyses have revealed that although there do not appear to be age differences in risky decisions made from description (in the absence of learning requirements), age differences in risk taking are more common on tasks where performance depends on learning from experience. Older adults also have lower levels of dopamine receptors and transporters in general (Karrer, Josef, Mata, Morris, & Samanez-Larkin, 2017). Studies have shown that dopamine receptor availability plays a role in reward learning deficits (de Boer et al., 2017) and that dopamine drugs can enhance reinforcement learning in older age (Chowdhury et al., 2013; Rutledge et al., 2009). It is not yet clear whether reduced dopamine in older age is associated with financial risk taking. Although neuroimaging studies of adulthood have shown dopamine reductions with age (Juarez et al., 2019) the same data also reveal no associations between dopamine and risky choice (Castrellon et al., 2019) using a task that does not depend on learning. However, learning-based risky decisions may be more strongly associated with individual differences in dopamine function across adulthood.

Along with scientific research in general, the reproducibility and generalizability of studies of brain-behavior individual difference effects have been critiqued and questioned recently (Dubois & Adolphs, 2016; Poldrack et al., 2017). This critique is well deserved. Many tasks produce multiple performance-related measures and whole-brain brain imaging always produces multiple measures. The experimenter degrees of freedom in brain-behavior studies are high in general and individual difference effects are regularly severely underpowered. This combination increases the likelihood of false positives or results produced completely from p-hacking. There is rarely a clear strategy for pairwise univariate analysis of data like those collected in the present study and many studies in this field. However, multivariate approaches are ideally suited for brain-behavior studies. Despite the ideal utility of these approaches, they are still rare. Even as multivariate neuroimaging studies increase, many of these studies use a single behavioral measure or limit the multivariate analyses to the brain imaging data. Yet, commonly the data are multivariate on both sides. Some emerging research has used data reduction techniques like partial-least squares to reduce dimensionality while evaluating brain-behavior associations (Calhoun, Liu, & Adalı, 2009; McIntosh & Lobaugh, 2004), but few studies have used techniques that combine multiple brain and behavioral measures in a single model. Multivariate methods may help better guide analysis of these naturally multivariate data sets and produce a more comprehensive understanding of brain-behavior associations in general.

The goal of this study was to examine multivariate associations between dopamine function and learning-based financial decision making in an adulthood sample. Using the [^11^C] FLB 457 radioligand, partial-volume-corrected (PVC) dopamine D2-like receptor non-displaceable binding potential (BP_ND_) was estimated from four regions of interest: midbrain, amygdala, anterior cingulate, insula. These regions of interest were identified based on previous research documenting critical contributions to learning (midbrain, amygdala) (O’Doherty, 2004) and risky decision making (anterior cingulate, insula) (Kuhnen & Knutson, 2005). Receptor availability cannot be reliably quantified in the striatum using [^11^C] FLB 457, otherwise this region would have also been included. Five measures of task-related errors during a financial investment task were computed to assess different aspects of learning-based decision making. To avoid excessive individual pairwise analyses given the number of variables on each side of the correlation, we used canonical correlation analysis (CCA; Hotelling, 1936) to identify associations between brain (BP_ND_ controlling for age) and behavior. We predicted that there would be an association between the set of brain variables and the set of task-related behavioral variables such that higher BP_ND_ would be associated with better performance (i.e., fewer task-related errors).

## Methods

### Participants

Thirty-seven healthy adults were recruited from the greater New Haven, CT community for a study of motivated cognition and decision making and written informed consent was obtained from all participants. Two participants were excluded (due to PET data quality issues), resulting in a final sample of 35 (ages 26-79 years; *M±*SD = 47.7*±*17.4). All participants completed extensive screening criteria, and thus were cognitively and physically healthy. Exclusion criteria included any history of psychiatric illness or head trauma, any significant medical condition, contraindications for MRI, history of substance abuse or psychostimulant use, and current use of tobacco, alcohol, or psychotropic drugs (see Castrellon et al., 2019, for a complete listing of exclusion criteria). All procedures and consent forms were approved by the Yale University Human Investigation Committee and the Yale-New Haven Hospital Radiation Safety committee.

### Risky Investment Decision Making Task

On each trial of the risky investment task, participants choose between two assets: a risky stock and certain bond (Figure 1, (Kuhnen, 2015)). Prior to beginning the task, participants were informed that there were two conditions, a gain condition where both assets would provide positive payoffs and a loss condition where both assets would provide negative payoffs. They were also informed that some stocks were *good*, and thus had a high probability of good outcome (winning or not losing money), while other stocks were *bad* and had a low probability of a good outcome (winning or not losing money) (see Appendix of Kuhnen, 2015, for complete instructions given to participants). Participants completed 10 practice trials before beginning the task. After the choice was made, the payoff for the stock was displayed, regardless of the choice made by the participant (to provide opportunities for learning independent of choice behavior) and then participants were then told their cumulative winnings. Finally, they were asked to estimate the probability that the stock is the “good” stock and their confidence in this probability estimate.

**Figure 1.**
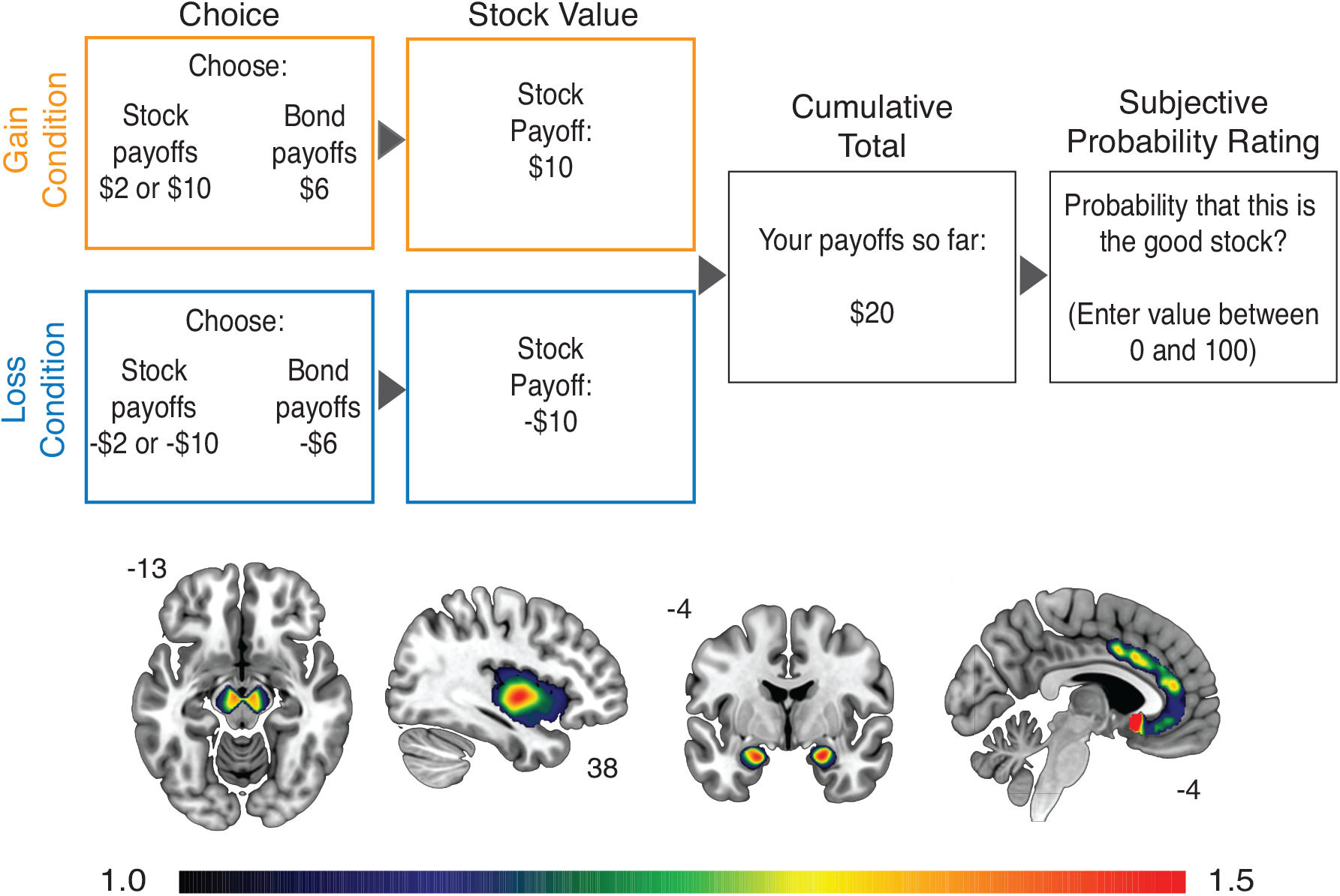
Task schematic for the learning-based risky investment decision task (top) and regions of interest (bottom) where dopamine receptor availability was extracted from the PET imaging data. The colorbar indicates levels of non-displaceable binding potential at the voxel level within these regions.

This was a 2 (framing: gain or loss) by 2 (stock payoff distribution: good or bad) design. In the gain condition, both assets provided positive payoffs (stock: +2 or +10; bond: +6) while in the loss condition, both assets provided negative payoffs (stock: -2 or -10; bond -6). Trials proceeded in 6-trial blocks. For each block, the feedback for stock selection was pulled from the same stock payoff distribution (*good* or *bad*). In the *good* stock payoff blocks, the high outcome (+10) occurs 70% of the time, while the low outcome (+2) occurs 30% of the time. In the *bad* stock payoff blocks these contingencies were flipped: the high outcome only occurs 30% of the time while the low outcome occurs 70% of the time. The order of framing conditions varied pseudorandomly and was the same for each participant. Participants completed a total of 10 blocks, for a total of 60 trials. The task was incentive-compatible such that participants were paid based on both their investment payoffs and accuracy in their probability judgments. Specifically, they received 1/10 of their cumulative payoffs plus 10 cents for each probability estimate within 5% of the objective Bayesian value in addition to hourly pay for participating in the study in general.

Task behavior was quantified using five error metrics: three measures of deviations from Bayesian reward maximization (choice inflexibility, first stock choice, and suboptimal investment) and two measures of probability estimation errors (absolute error and optimism). The Bayesian reward maximization metrics were calculated based the stock vs bond choices made by participants. Choice inflexibility quantifies how much the participant persisted in choosing the asset that they chose in the first trial of the block, when this asset was objectively the less optimal choice. First stock choice, which is a measure of risk-taking propensity, is the proportion of the time that participants chose the stock (versus the bond) on the first trial of a block. Suboptimal investment is the proportion of trials where the participant chose the wrong asset. The probability estimation errors were based on the “good-stock” follow-up questions. Optimism is the amount a person overestimates the probability that they have chosen the “good stock”, or more precisely it is the average distance between a participant’s probability estimate and the objective Bayesian posterior probability estimate, which is

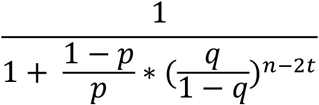

for *n* trials so far and *t* high outcomes so far when the prior that the stock is “good”, *p =* 50%, the probability that the good stock pays the high (compared the low) dividend in each trial, *q* = 70%. Absolute error is simply the absolute value of optimism. It quantifies how inaccurate the participant’s probability estimates are in either direction, whether they are overestimating or underestimating the probability of a stock being the “good stock”.

### PET data acquisition and processing

PET imaging was collected at Yale New-Haven Medical Center. [^11^C] FLB 457, 5-bromo-N-[[(2S)-1-ethyl-2-pyrrolidinyl]methyl]-3-methoxy-2-(methoxy-11C) benzamide was synthesized in the Yale PET Center radiochemistry laboratory in the Yale School of Medicine. PET scans were acquired on a high resolution research tomograph (HRRT; Siemens Medical Solutions, Knoxville, TN, USA). [^11^C] FLB 457 (median specific activity: 7.80 mCi/nmol) was injected intravenously as a bolus (315 MBq; average 8.62 mCi, SD 2.03 mCi) over 1 min by an automated infusion pump (Harvard Apparatus, Holliston, MA, USA). Prior to each scan, a six-minute transmission scan was performed for attenuation correction. Dynamic scan data were acquired in list mode for 90min following the administration of [11C]FLB 457 and reconstructed into 27 frames (6×0.5 mins, 3×1min, 2×2mins, 16×5 mins) with corrections for attenuation, normalization, scatter, randoms, and dead time using the MOLAR (Motion-compensation OSEM List-mode Algorithm for Resolution-Recovery Reconstruction) algorithm. Event-by-event, motion correction was applied using a Polaris Vicra optical tracking system (NDI Systems, Waterloo, Canada) that detects motion using reflectors mounted on a cap worn by the subject throughout the duration of the scan. After decay correction and attenuation correction, PET scan frames were corrected for motion using SPM8 (Friston et al., 1994) with the 13th dynamic image frame of the first series serving as the reference image. The realigned PET frames were then merged and reassociated with their acquisition timing info in PMOD’s PVIEW module to create a single 4D file for use in PMOD’s PNEURO tool for further analysis.

### MRI data acquisition

Structural MRI scans were collected using a Siemens Trio whole-body scanner (Siemens Medical Systems, Erlangen, Germany). T1-weighted high-resolution anatomical scans (repetition time = 2.4 s, echo time = 1.9 ms, field of view = 256 x 256, voxel dimensions = 1 x 1 x 1 mm) were obtained for each participant. These structural scans facilitated co-registration and spatial normalization of the PET data.

### Partial volume correction and brain regions of interest

Both MRI and PET data were parcellated into 62 bilateral cortical, 12 bilateral subcortical, 3 posterior fossa, 5 ventricle, and 1 white matter regions of interest (a total of 83 regions) using the Hammers atlas (Gousias et al., 2008; Hammers et al., 2003). Following parcellation, the MRI and PET data were co-registered, PET data was resampled to MRI space, and then the partial volume correction (PVC) procedure available in PMOD’s PNEURO module was applied to the PET data. PNEURO uses the GTM method (Rousset, Collins, Rahmim, & Wong, 2008; Rousset, Ma, & Evans, 1998), which restricts PVC to the PET signal of structurally defined regions of interest. After PVC, time activity curves (TACs) from each region were extracted from the PET data and fit with a simplified reference tissue model (Lammertsma & Hume, 1996) where a gray matter bilateral cerebellum ROI was used as the reference region using PMOD’s PKIN module (see Smith et al., 2019), for greater detail). Four regions of interest (ROIs) were selected from the Hammers atlas *a priori* for analysis: midbrain, amygdala, insula and anterior cingulate. Within each participant, mean non-displaceable binding potential (BP_ND_) was calculated for each of these ROIs (Figure 1).

### Canonical Correlation Analysis

To test for an association between dopamine receptor availability measures and decision-making task measures we performed a canonical correlation analysis (CCA). CCA is a general statistical method that measures the relationship between two multidimensional variable sets. More specifically, given two datasets X and Y, CCA seeks to find a linear combination of X that maximally correlates with a linear combination of Y. The linear combinations of X and Y are synthetic variables, called canonical variates. The canonical variates together are a canonical pair and the correlation between the canonical variates is called a canonical correlation. The possible number of canonical pairs is limited to the number of variables in the smallest dataset with the constraint that each successive pair is uncorrelated with all previous ones.

Statistical significance was defined at p < 0.05. Analysis was conducted in R (Version 3.6.1; R Core Team, 2019) using the CCP (Version 1.1; Menzel, 2012) and the yacca (Version 1.1.1; Butts, 2018) packages.

## Results

A canonical correlation analysis (CCA) was conducted using four dopamine D2-like receptor availability variables (midbrain, amygdala, insula, anterior cingulate) and age as correlates of the five financial decision making task performance variables (choice inflexibility, risk seeking, suboptimal investment, absolute error, and optimism). The CCA evaluated the multivariate shared relationship between the two variable sets (i.e. dopamine receptor availability controlling for age and risky investment decision making). The analysis produced five functions with squared canonical correlations (R^2^_c_) of .581, .351, .072, .028, and .005. The full model across all five functions was statistically significant using the Wilks’s = .24 test criterion, *F*(25, 94.373) = 1.747 *p* = .029. The effect size for the complete set of five canonical correlations was *r*^*2*^ *=* .581, showing that the full model explained 58% of the shared variance between the two variable sets. A dimension reduction analysis showed that the second through fifth functions did not explain statistically significant amounts of shared variance as seen in Table 1. As such, the following results are focused on the first function (see Figure 2).

**Table 1.**
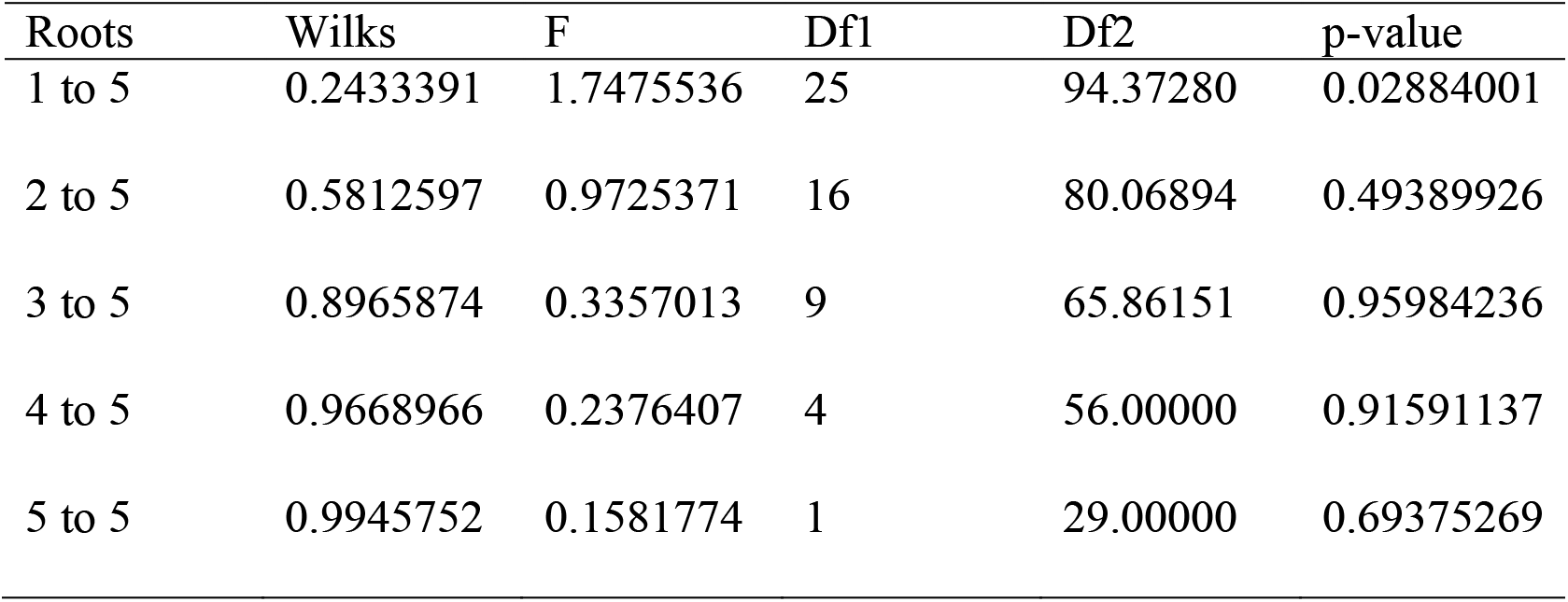
Dimension reduction showing the significant shared variance overall but not in the second through fifth functions.

**Figure 2.**
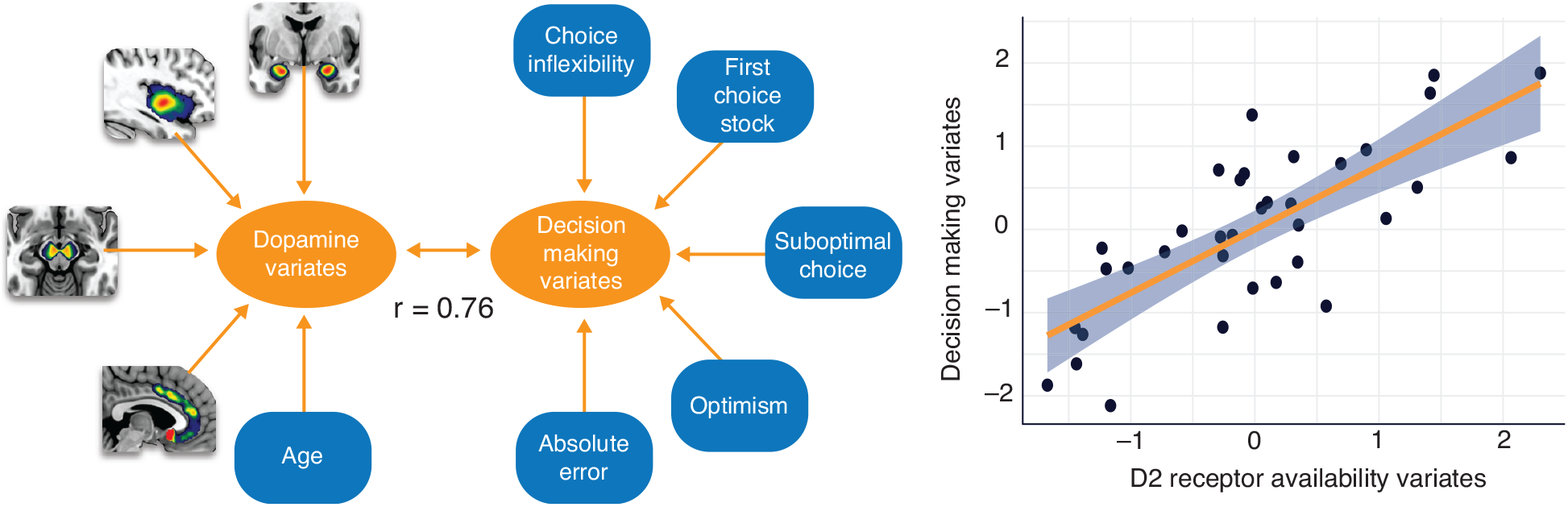
Dopamine receptor availability was associated with task performance overall (r = 0.76).

Table 2 shows the standardized canonical coefficients and structure coefficients for Function 1, along with the squared structure coefficients. The structure coefficients for the criterion variables suggest that choice inflexibility, incorrect choice, and optimism are the relevant contributors to the decision making variate. Each of the criterion variables also had large standardized canonical function coefficients, except for incorrect choice which only had a moderate coefficient due to multicollinearity with the other decision-making variables. The structure coefficients for inflexibility and incorrect choice have the same sign, indicating that they were positively related. Optimism was inversely related to the other criterion variables.

**Table 2.** Standardized canonical coefficients (Coef), structure coefficients (r_s_), and squared structure coefficients (*r*_*s*_^*2*^) for each variable for Function 1.

| | *Coef* | $r_s$ | $r_s^2$ |
| --- | --- | --- | --- |
| *Age and brain variables* |  |  |  |
| Age | 0.06 | 0.61 | 0.37 |
| Midbrain | 2.27 | -0.14 | 0.02 |
| Amygdala | -1.03 | -0.52 | 0.27 |
| Insula | -0.72 | -0.37 | 0.14 |
| Anterior cingulate | 1.30 | -0.08 | 0.01 |
| *Task variables* |  |  |  |
| Choice inflexibility | 7.76 | 0.72 | 0.52 |
| Chose stock first | 1.29 | 0.35 | 0.12 |
| Suboptimal investment | 0.96 | 0.57 | 0.33 |
| Probability estimation error | .003 | 0.33 | 0.11 |
| Optimism | -0.05 | -0.49 | 0.24 |

From the predictor variables, structural coefficients suggest that age and amygdala dopamine binding potential were the relevant contributors to the predictor variate. Age was positively related to each of the relevant task-related variables except for optimism. Amygdala binding potential was negatively related to all the task-related variables except for optimism.

Follow-up exploratory univariate analyses were conducted to examine the effects of dopamine receptor availability within individual ROIs and age on choice inflexibility using more traditional analysis methods (see Figure 3). Univariate linear regression showed a significant positive association between age and choice inflexibility b = .002, t(33)=2.44, p = 0.02, and negative association between dopamine receptor availability within the amygdala and choice inflexibility b = –0.04, t(33) = –2.15, p = 0.04, controlling for age. This negative correlation indicated that higher levels of dopamine receptor availability were associated with better performance (less inflexible or more flexible choice). Age explained a significant proportion of variance in choice inflexibility, R^2^=0.15, p =.02. Amygdala dopamine receptor availability also explained a significant proportion of variance in choice inflexibility, R^2^=0.12, p =.03.

**Figure 3.**
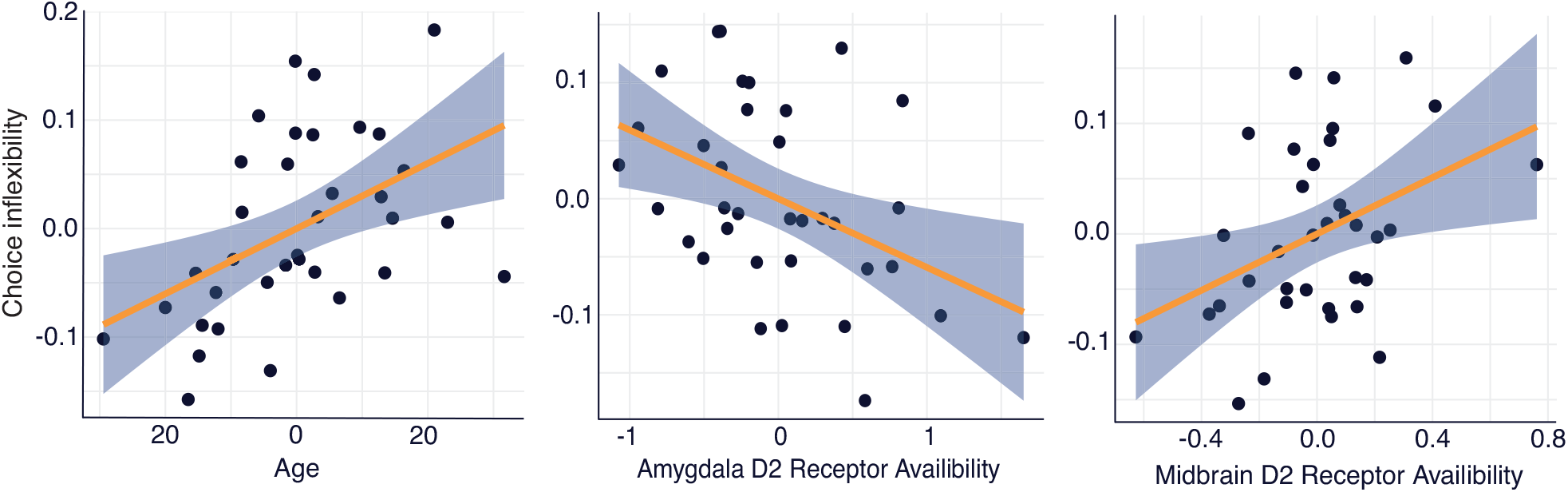
Univariate associations between age and choice inflexibility, amygdala dopamine receptor availability and choice inflexibility, and midbrain dopamine receptor availability and choice inflexibility.

## Discussion

The present study evaluated whether baseline dopamine receptor availability was related to individual differences in learning-based risky financial decision making across adulthood. In a relatively small sample of healthy human adults, we identified preliminary multivariate associations between risky investment decision making and dopamine receptor availability. Using multivariate CCA, the study reveals general associations between dopamine receptor availability and risky decision making. In light of the mixed suggestions from pharmacological studies about the effects of dopamine on risky behavior, the present data suggest that baseline levels of dopamine across adulthood are directly related to risky financial decision making. Thus, it is important to understand an individual’s baseline dopamine status when evaluating how manipulations of the dopamine system might affect decision making.

Decomposition of the CCA and follow-up univariate analyses demonstrated that the effects were driven by negative correlations between amygdala D2-like receptor availability and choice inflexibility controlling for age. Individuals with lower levels of D2-like receptor availability in the amygdala were more likely to persist in choosing the asset that they chose in the first trial of the block when this asset was objectively the less optimal choice. Individuals with higher levels of D2-like receptor availability in the amygdala were less likely to make these perseverative errors. Functional neuroimaging studies have identified contributions of amygdala function to risky decision making in general (Bechara, Damasio, & Damasio, 2003; Kuhnen & Knutson, 2005) as well as individual difference associations between neural activity in the amygdala and risky decision making (De Martino, 2006). The current results provide new suggestive evidence that some portion of these effects might be driven specifically by dopamine receptors within the amygdala. Although neuroimaging research on financial decision making has focused more heavily on corticostriatal dopamine circuits, there are prominent anatomical dopamine projections from the midbrain to the medial temporal lobes (Berger, Gaspar, & Verney, 1991) and specifically into the amygdala. Prior PET imaging studies show relatively high levels of dopamine D2-like receptor availability in the amygdala (Okubo et al., 2000).

The effects in this study are novel in both the specific implication of dopamine function in the amygdala and the specific contributions of learning to risky choice. Individuals with low amygdala D2 receptor availability seem to be less sensitive to feedback when their initial choice ends up being a worse option. In this phase of the task, they would be better off choosing the safe asset, the bond, but instead persist in choosing a losing stock. Although this task is inspired by investment decision making in everyday life, we did not collect measures of individuals real-life reactions or choices after negative events in the stock market for example. The task was incentive-compatible but the financial stakes were relatively small. Although suggestive, it’s unclear whether these effects would extend to perseveration on poor initial choices in everyday risk taking.

The associations between amygdala receptor availability and choice inflexibility held controlling for age in univariate analyses. However, age was also correlated with choice inflexibility. The strength of the independent associations between choice inflexibility and both age and amygdala receptor availability were similar with each accounting for 12-15% of the variance in perseveration on the first choice made in a block after trial-related feedback revealed that it was a bad choice. These effects suggest that adults may be less likely to adjust their choice behavior after receiving feedback that they are losing money. The suggestion that older adults might be more likely to make perseverative errors in investment decision making would be consistent with a wealth of research on aging and perseveration in general (Hartman, Bolton, & Fehnel, 2001; Hosseini et al., 2010). Although age is negatively correlated with dopamine receptor availability in nearly all regions of the brain, the effects of age on dopamine D2-like receptors in the amygdala are relatively small (Seaman et al., 2019). That is, dopamine receptor availability in this region appears to be relatively preserved with age in general. Together the effects suggests that the older adults with the lowest levels of amygdala D2 receptor availability may be the most likely to perseverate on losing options in investment decision making. Critically, the sample size in this study is small so it is not possible to generalize much from the observed age effects overall or specifically make generalizations about individual difference effects within age groups. Much more data would be required to evaluate potential interactions between age and amygdala dopamine receptor availability.

In summary, the study provides novel evidence for multivariate associations between dopamine receptors and risky learning-based investment decision making across adulthood. Given the small sample size, the replicability and generalizability of the effects is not yet clear. However, the study does provide suggestive evidence for the utility of using a multivariate approach to examining brain-behavior associations in adulthood. Given how frequently behavioral tasks can produce multiple performance-related measures (e.g., speed, accuracy) and the number of data points collected in typical brain imaging studies, multivariate methods are ideally suited for identifying brain-behavior associations. Discussions of questionable research practices have regularly identified experimenter degrees of freedom as a potential source of bias in the literature (Wicherts et al., 2016). Initial omnibus statistical testing with multivariate tools may help better organize and identify overall patterns of association across sets of measures for more strategic decomposition of effects. Of course, more data than has typically been collected (Button et al., 2013) will be required for evaluation of these multivariate effects assuming the true effects are relatively small (Dubois & Adolphs, 2016). However, so little research has used multivariate brain-behavior association approaches that the size of the true effects is currently unknown in general and especially for any specific set of behavioral or brain measures. This is the first study of which we are aware that examined multiple measures of dopamine function and decision-making behavior in a single model. Other emerging research has used related multivariate approach with dopamine imaging data in humans but those studies have been focused on more traditional cognitive measures like memory (Guitart-Masip et al., 2016; Lövdén et al., 2018; Salami et al., 2019). As sample sizes increase, these multivariate approaches may prove more generally useful in understanding variation across individuals in brain function and the effects on other aspects of behavior.

## Acknowledgements

This study was supported by National Institute on Aging grant R00-AG042596 to G.R.S.-L. and T32-AG000029 to K.L.S..

